# The rate of spontaneous mutations in yeast deficient for MutSβ function

**DOI:** 10.1101/2022.08.25.505291

**Authors:** Yevgeniy Plavskin, Maria Stella de Biase, Roland F Schwarz, Mark L. Siegal

## Abstract

Mutations in simple sequence repeat loci underlie many inherited disorders in humans, and are increasingly recognized as important determinants of natural phenotypic variation. In eukaryotes, mutations in these sequences are primarily repaired by the MutSβ mismatch repair complex. To better understand the role of this complex in mismatch repair and the determinants of simple sequence repeat mutation predisposition, we performed mutation accumulation in yeast strains with abrogated MutSβ function. We demonstrate that mutations in simple sequence repeat loci in the absence of mismatch repair are primarily deletions. We also show that mutations accumulate at drastically different rates in short (<8 bp) and longer repeat loci. These data lend support to a model in which the mismatch repair complex is responsible for repair primarily in longer simple sequence repeats.

## Introduction

Simple sequence repeats (SSRs) are consecutive repeats of short nucleotide sequences found throughout genomes. Their arrangement makes SSRs particularly susceptible to mutation (Strand *et al*. 1993). SSR variants are increasingly recognized as contributing to natural phenotypic diversity, including variation in gene expression in humans (Gymrek *et al*. 2016; Fotsing *et al*. 2019) and yeast (Vinces *et al*. 2009), with potential consequences for the evolution of promoters (Sonay *et al*. 2015). Moreover, mutations in SSR loci play a key role in many inherited human health conditions: both Huntington disease and Fragile X syndrome are caused by expansions of triplet repeats, as are tens of other human disorders (reviewed in López Castel *et al*. 2010).

SSR mutations are caused by polymerase slippage events during replication, and in most cases are normally corrected by the cell’s mismatch repair (MMR) mechanism (Strand *et al*. 1993, 1995). In eukaryotes, mismatches are recognized and repaired by one of two heterodimers: MutSα, consisting of Msh2p and Msh6p, which primarily repairs mismatches and single-nucleotide insertions and deletions, and MutSβ, consisting of Msh2p and Msh3p, which primarily repairs short insertions and deletions (reviewed in (Kunkel and Erie 2015)), although mutations in *MSH3* also result in changes in the spectrum of substitutions (Harrington and Kolodner 2007; Lamb *et al*. 2022). Mutations in the MMR complexes, especially MutSα, play a key role in certain cancers, and are the primary genetic basis of Lynch syndrome, which results in a high rate of colorectal, endometrial, and other cancers (Heinen 2016). Natural variants in *MSH3* also influence the rate of SSR germline mutations in mice (Maksimov *et al*. 2022) and the severity and age of onset of two diseases caused by SSR expansions in humans, Huntington Disease and Myotonic Dystrophy Type 1 (Flower *et al*. 2019).

Many studies have examined the role of MMR complexes in SSR mutation repair using reporter constructs in which an SSR precedes a reporter gene in yeast, such that SSR indels result in a selectable phenotypic readout (e.g. (Strand *et al*. 1993, 1995) and others). By combining this reporter with MMR mutants, the relative roles of the various MMR components in indel repair have been elucidated. This work has demonstrated biases in insertion/deletion rates of new mutations, the dependence of mutation rate on SSR length, and in some cases, higher mutation rates in G/C-rich SSRs (Kunkel and Erie 2015). One key strength of the reporter gene approach is that individuals in which the reporter SSR locus is mutated can be easily identified and sequenced in a targeted manner, and newer protocols have allowed increased throughput of this approach (Ollodart *et al*. 2021). However, such studies are limited to studying a single artificial SSR locus at a time. Recent work has instead leveraged high-throughput sequencing of mutation accumulation lines to understand the dependence of SSR mutation rate on both mismatch repair and SSR locus properties (Lynch *et al*. 2008; Lang *et al*. 2013; Serero *et al*. 2014; Lujan *et al*. 2015; Haye and Gammie 2015; Belfield *et al*. 2018; Saxena *et al*. 2019; Konrad *et al*. 2019). In these studies, individuals are propagated with extreme bottlenecks over many generations, resulting in fixation of mutations primarily by drift, largely unaffected by their phenotypic effect. At the end of the mutation accumulation, strains are sequenced, and mutations in SSR loci are identified genome-wide. Such studies have led to estimates of SSR mutation rate in wildtype *C. elegans* (Saxena *et al*. 2019; Konrad *et al*. 2019) and yeast (Lynch *et al*. 2008), MMR-deficient yeast (Lang *et al*. 2013; Lujan *et al*. 2014, 2015; Haye and Gammie 2015) and *Arabidopsis* (Belfield *et al*. 2018), as well as in yeast with defects in polymerase proofreading (Lujan *et al*. 2015). These studies confirm many of the properties of SSRs that confer increased mutation rates first identified in reporter studies. Furthermore, Lujan *et al*. (2015) showed that indels in SSRs <5-10 nucleotides long had increased rates of mutation when both MMR and polymerase proofreading were abrogated, suggesting that indels in short SSRs are repaired by both the MMR complex and polymerase, whereas mutations in longer SSRs are repaired primarily via MMR.

Although mutation accumulation-based studies have been able to assess the mutation rates of SSR loci at genome-wide scale, a persistent challenge lies in the difficulty of accurately calling SSR alleles with high-throughput sequencing data (Gymrek *et al*. 2012). Genotyping error rates in SSR loci are correlated with *in vivo* mutation rates, making it more difficult to disentangle genotyping error rates from differences in mutation rates across loci, especially when the total mutation number is low. Furthermore, most genome-wide studies of MMR function have been performed in strains in which either both MutS complexes, or only MutSα, were perturbed.

To better understand the role of MutSβ in repair of mutations in SSR loci, we performed a mutation accumulation experiment in *Saccharomyces cerevisiae* strains in which the *MSH3* gene was deleted, leaving the MutSα complex intact. We accumulated mutations over 200 generations in more than 30 haploid lines. To avoid bias in mutation rate estimation, we analyzed mutation rates by setting stringent calling quality thresholds within groups of SSR loci with related properties. We show that this method results in consistent estimates of the overall SSR mutation rate across a wide range of thresholds. We find that the loss of *MSH3* results in an increase in the SSR mutation rate, but do not observe an increase in the rate of single-nucleotide mutations (SNMs) or non-SSR indels. We also find that in strains with an *msh3* deletion (*msh3Δ*), per-basepair mutation rate is much higher in long SSRs. Finally, we confirm the predominance of SSR deletion mutations in *msh3Δ* mutation accumulation strains.

## Results

### *Deletion of* MSH3 *does not significantly affect single-nucleotide mutation rate outside of simple sequence repeats*

Previous work indicates that Msh3p, and the MutSβ complex it is part of, is a key component of the mismatch repair system for small indels, but not for substitutions (reviewed in Kunkel and Erie 2015). To test whether deletion of *MSH3* affected substitution rate, we performed mutation accumulation (MA) for ∼200 generations on wildtype and *msh3Δ* haploid strains (**Figure 1**). We sequenced five wildtype MA lines and 34 *msh3Δ* MA lines, as well as the MA ancestors (**Supplementary Table 1**). Mutations were called with both FreeBayes (Garrison and Marth 2012) and muver (Burkholder *et al*. 2018) to increase call set completeness. muver calls mutations relative to a sample specified as the ancestral strain. In FreeBayes, alleles were called relative to the reference sequence and alleles that differed between ancestral and MA lines were called as mutations in the MA lines (see Materials and Methods). We filtered all FreeBayes-identified mutations to select mutations sequenced at at least 10x coverage and outside of SSR regions. All muver-identified mutations outside of SSR regions, except two, were a subset of the filtered mutations identified by FreeBayes; Sanger sequencing revealed that these two mutations were false positives. We therefore performed further analysis using FreeBayes mutation calls.

**Figure 1:**
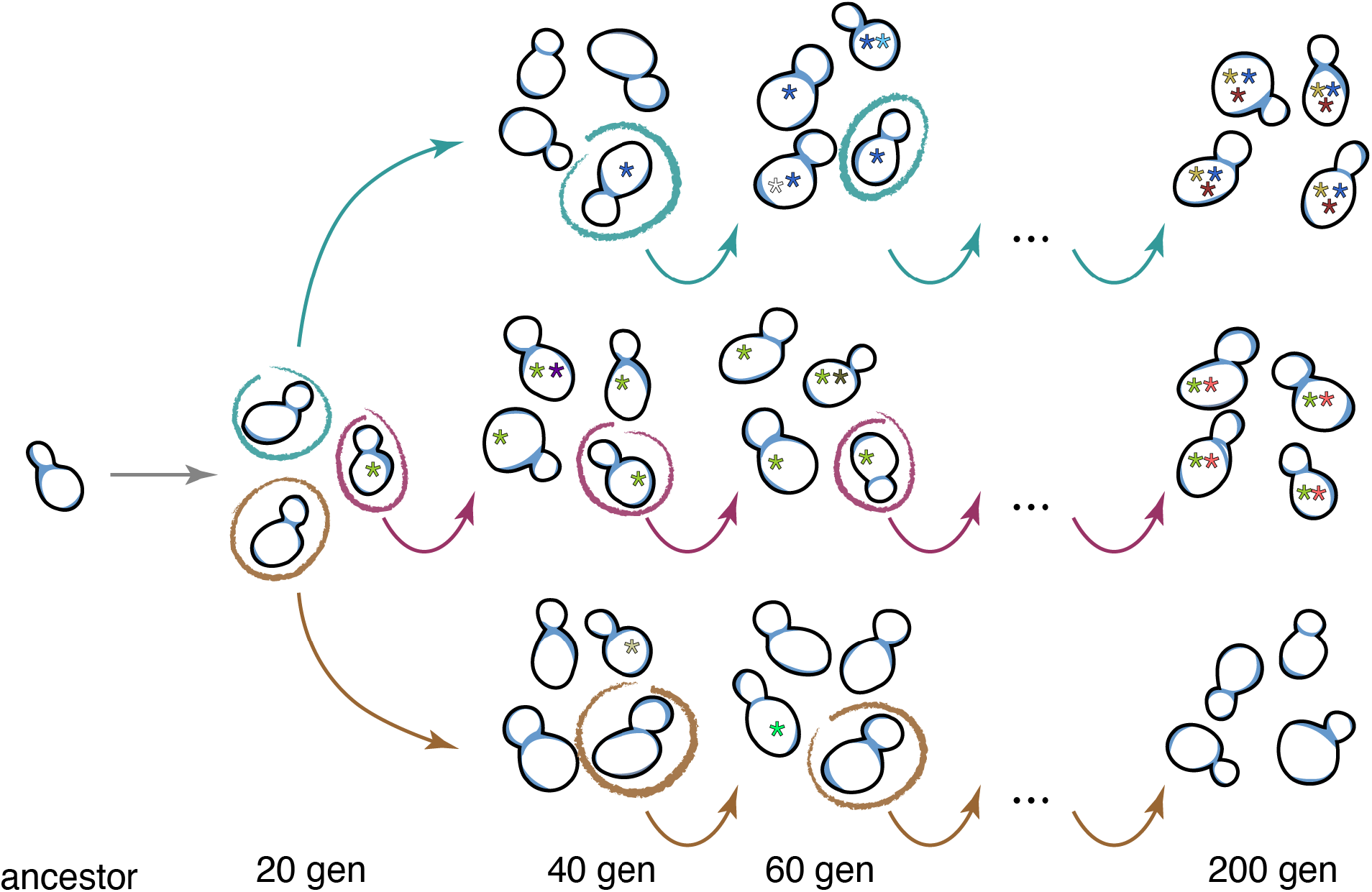
Mutation accumulation scheme. Multiple lines were started from a single colony derived from one ancestral cell, and each line was propagated through bottlenecks of a single, randomly chosen cell every ∼20 generations by re-streaking a single colony, for a total of 10 transfers. Mutations (asterisks) randomly accumulated in the cells over the course of the experiment.

24 substitutions passed filtration in multiple MA strains, together accounting for 213 mutation calls (85% of all filtration-passing substitution calls). Because all MA lines are independently derived from a single common ancestor, the chance of two or more lines sharing a mutation is very low. An additional 10 substitutions were found a small distance (<50 bp away) from another substitution in a different strain, also highly unlikely to occur by chance considering that across all strains, only 27 unique substitutions were called elsewhere in the genome. We therefore further examined these calls. We first selected single-nucleotide substitution mutations, grouped mutations found within 50-bp of each other, and determined the number of strains that had a mutation within each such locus. This allowed us to identify regions with recurring mutation calls across strains. We found that loci mutated across multiple strains had properties distinct from single-strain substitutions: unlike uniquely mutated loci, all loci with mutations in multiple strains had, in either the MA line or its ancestor, a mix of reads supporting two different alleles, with the proportion of reads supporting the not-called allele being >25% of the total mapped reads. Strains without a called mutation at these loci also had a high proportion of reads supporting alleles other than the called allele; this was not the case for filtration-passing loci at which a single strain was mutated (**Supplementary Figure 1**). Together, these results suggest that mutations detected in multiple strains are the result of unreliable calls. We therefore removed any loci with SNMs mutated in multiple strains from our analysis.

The final FreeBayes mutation call-set consisted of 27 SNMs outside SSR regions, sequenced at >10x coverage and present in a single strain **(Supplementary Tables 2-3)**. We checked all of these mutations using Sanger sequencing, which confirmed all 27 identified mutant loci. Of these loci, 2 were in a single *MSH3*^*+*^ strain, and the rest were found among 17 of the 34 sequenced *msh3*Δ strains. To compute the non-SSR SNM mutation rate across strains, we considered the non-repetitive proportion of the genome sequenced at 10x or higher. We found no significant difference between the non-SNM mutation rate in *MSH3*^*+*^ and *msh3*Δ MA strains (p = 0.36); the non-SSR SNM mutation rate across all strains is 0.8 mutations/genome (over the course of ∼200 generations), or 3.5 × 10^−10^ mutations/bp/generation (95% CI: 2.3-4.9× 10^−10^ mutations/bp/generation).

### Property-dependent filtering strategy for SSR mutation calls reduces bias against highly mutable loci

Using the same filtration parameters as with SNMs, we identified and confirmed by Sanger sequencing five insertions and deletions outside of identified SSR regions, as defined in Materials and Methods. However, closer inspection revealed that all of these occurred in SSRs that were not labeled as such by our original algorithm (**Supplementary Table 4**), either because the motif length was >4 nucleotides, or because the unmutated SSR had too few perfect repeats of the motif to be recognized.

Unlike SNMs, applying the same call-confidence-based cutoff to mutations within SSRs is likely to produce incorrect estimates of the SSR mutation rate. PCR introduces stutter noise into reads containing SSR loci (Willems *et al*. 2017); as a result, multiple reads often contain identical insertions or deletions by chance, potentially requiring a different likelihood cutoff to be used for SSR loci than for SNMs in the rest of the genome. Furthermore, we hypothesized that the likelihood assigned to a call may be negatively correlated with the probability of a mutation occurring at a locus. There are two potential reasons for this. First, because the mechanism of SSR locus mutation is polymerase slippage (Strand *et al*. 1993), the probability of PCR stutter at a locus may be correlated with the probability of that locus being mutated *in vivo*, introducing increased uncertainty in calls at loci that are more likely to be mutated. Second, accurate calls of SSR locus length require reads to completely span the locus. Longer loci, which are more likely to be mutated, are likely to have fewer spanning reads and thus have lower call confidence. Indeed, we find that the distribution of likelihood differences between the top two most likely allele calls (ΔGL) at an SSR locus varies depending on key properties such as SSR motif length, total repeat length, and A/T proportion (**Figure 2A, Supplementary Figure 2A**), all factors previously shown to be correlated with SSR mutation rate (e.g. Lang *et al*. 2013; Lujan *et al*. 2015).

**Figure 2:**
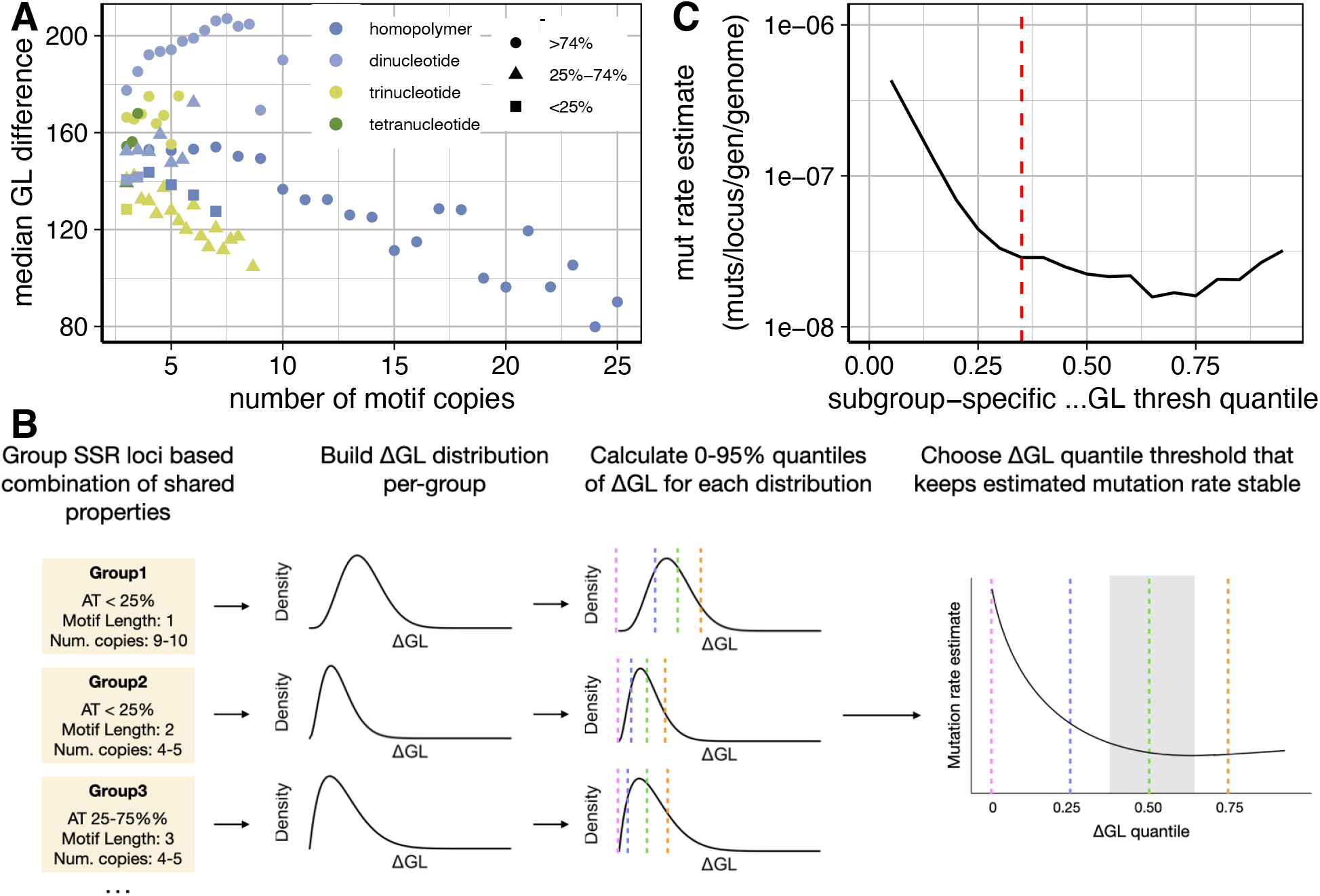
Property-dependent filtering of SSR locus calls. (A) Median call confidence (ΔGL; difference in log10(phred-scaled genotype likelihood) between top two most likely alleles) increases with SSR motif complexity and A/T proportion, and decreases with number of motif copies. Plot includes only groups containing >25 distinct SSR loci. (B) Scheme of call filtration. Similar SSR loci are grouped, and the number of mutant/non-mutant loci is calculated at various quantiles of ΔGL values; the mutation rate across all loci is then calculated for each quantile; the appropriate ΔGL threshold is chosen within the range of quantiles that maintains the calculated SSR mutation rate stable (gray shaded area); see also **Supplementary Figure 2** for examples of empirical ΔGL distributions. (C) Filtering using a minimum ΔGL difference threshold corresponding to a specific quantile of ΔGL differences within each group produces a stable estimate of SSR locus mutation rate across a wide range of threshold quantile values for haploid MA lines; red line represents the cutoff used in this study (35%).

The possible correlation of SSR mutation call confidence with SSR mutation rate means that using a single confidence-based cutoff to filter calls at SSR loci will bias any assessments of factors contributing to SSR mutation rate: the filter will be inherently less stringent for loci less likely to be mutated, resulting in a higher perceived mutation rate in those loci. To avoid this bias, we set up a custom filtration strategy, in which the appropriate filtering thresholds are applied to individual groups of SSR loci with matching properties (**Figure 2B, Supplementary Figure 2A**). We grouped all FreeBayes’ calls at loci with matching motif length, similar A/T proportion (<25%, 25%-75%, >75%), and similar motif copy number, and set up filtering thresholds based on the distribution of ΔGL values for each group of SSR calls. To avoid SSR groups with a small number of calls, for each unique motif copy number we considered a window of -2.5 to +2.5 repeats around the actual copy number (choice of motif copy number window size did not affect the results; **Supplementary Figure 2B**). To avoid spuriously low thresholds in groups of poorly called loci, we also required that each threshold be based on the ΔGL value of a group of calls including >25 individual SSR loci; loci in smaller groups were removed. Despite this, we were still able to make calls at many loci across different motif lengths, A/T proportion values, and repeat copy numbers (**Supplementary Figure 3B**). This strategy tailors the ΔGL threshold applied to each group of SSR loci to the distribution of ΔGL values observed across similar loci, accounting for differences in call confidence between loci with diverse properties. Importantly, these thresholds are then applied to all SSR calls in a mutation-agnostic manner (see Materials and Methods), so that both putatively mutated and wild-type loci are thresholded in the same way.

We tested quantile thresholds corresponding to the removal of anywhere from 0 to 95% of all allele calls. Because calls were made at nearly all SSR loci, regardless of whether they were mutated, we can estimate the proportion of mutant loci at each threshold, by merging the calls passing the chosen threshold in each group of SSR loci (**Figure 2C**). Because incorrect calls are mostly false positives (as the overall mutation rate is low), low filtration thresholds result in an inflated apparent mutation rate, with increasingly stringent thresholds decreasing the number of false positives and thus the apparent mutation rate. However, the measured SSR mutation rate is stable for thresholds corresponding to removal of 35-60% of all calls. At more stringent filtration thresholds, low overall mutation numbers result in large fluctuations to the mutation rate. Our filtration method is therefore not sensitive to precise thresholding parameters, and allows us to call SSR mutations without a bias against frequently mutated loci. We thus chose to remove the 35% of calls with the lowest ΔGL values in each SSR group. Although a cutoff anywhere in the 35-60% range would be suitable for determining the overall mutation rate, more stringent cutoffs remove more true positive mutations and true negative SSRs from our dataset. Using the cutoff at the low end of the stable range (35%) preserves more called loci, increasing statistical power in subsequent analyses.

We also compared the calls that passed filtration in our MA lines to calls made by muver within SSR regions and by MSIsensor, a program for calling SSR mutations in mutated samples relative to a control sample (Niu *et al*. 2014; Jia *et al*. 2020). MSIsensor detects microsatellite instability by comparing the distribution of reads corresponding to different motif numbers at each locus in a sample to a ‘normal’ control (in this case, the MA ancestor of each strain), although it excludes any reads that are not perfect repeats of the SSR motif (e.g. those containing indels different from the motif length or substitutions). Although our goal was not to call every mutation, but rather to correctly estimate the proportion of loci at which mutations occurred, we reasoned that the loci we identify with mutations should be a subset of a less stringently filtered set of loci. Consistent with this reasoning, 30 of 90 mutations identified by muver and 27 of 59 mutations identified by MSIsensor were also called in our analysis. We also identified a further 9 mutations not called by MSIsensor and 6 mutations not called by muver (2 of which were not called by both tools). These differences might be due to the different methods and filtering steps of the different tools. Indeed, all of the 9 mutations not called by MSIsensor were at loci that were excluded by MSIsensor’s algorithm due to technical reasons (e.g. imperfect repeat), and 2 of the mutations not identified by muver were in regions filtered out because of insufficient coverage according to muver’s algorithm.

### *Mutation rate at SSR loci is influenced by* MSH3 *status and locus length*

After filtering mutations as described above (removing 35% of calls with the lowest ΔGL values in their categories), we detect mutations in a total of 36 SSR loci (**Supplementary Table 5**), with 0-5 SSR loci mutated per strain (median mutation number = 0). The proportion of SSRs that are called also varies significantly among strains, likely due to variable average read depth among strains, with a median of 63% of all SSR loci called.

We first sought to characterize the determinants of the SSR mutation rate in *msh3*Δ strains. To do this, we modeled the odds of a mutation in an SSR as a per-base pair odds of mutation multiplied by the length of the SSR; in turn, the per-base pair mutation odds are modeled as the product of coefficients corresponding to various properties of the SSR and the strain’s *MSH3* status. We do not include the 5 indels identified in unannotated SSR loci in this modeling; because we do not have a complete list of loci with similar properties (e.g. with motif size >4) we are not able to apply the same filtering criteria as for the other SSR mutations, which could lead to incorrect mutation rate estimate.

Previous work has shown that the proportion of A/T nucleotides in the SSR influences the SSR mutation rate in *msh2* mutant strains (Gragg *et al*. 2002); A/T proportion also influences the error rate in SSR calls, likely due to altering the rate of the introduction of indels during PCR (**Figure 2A**). Among the SSR mutations in our *msh3*Δ MA strains, G/C-rich SSR loci appear overrepresented relative to A/T rich loci; however, the effect of A/T proportion on the SSR mutation rate is not statistically significant (*p* = 0.25). We also do not find a significant effect of motif length (homopolymer, dimer, or trimer, tetramer) on mutation rate (p = 0.5) (**Table 1**).

**Table 1:**
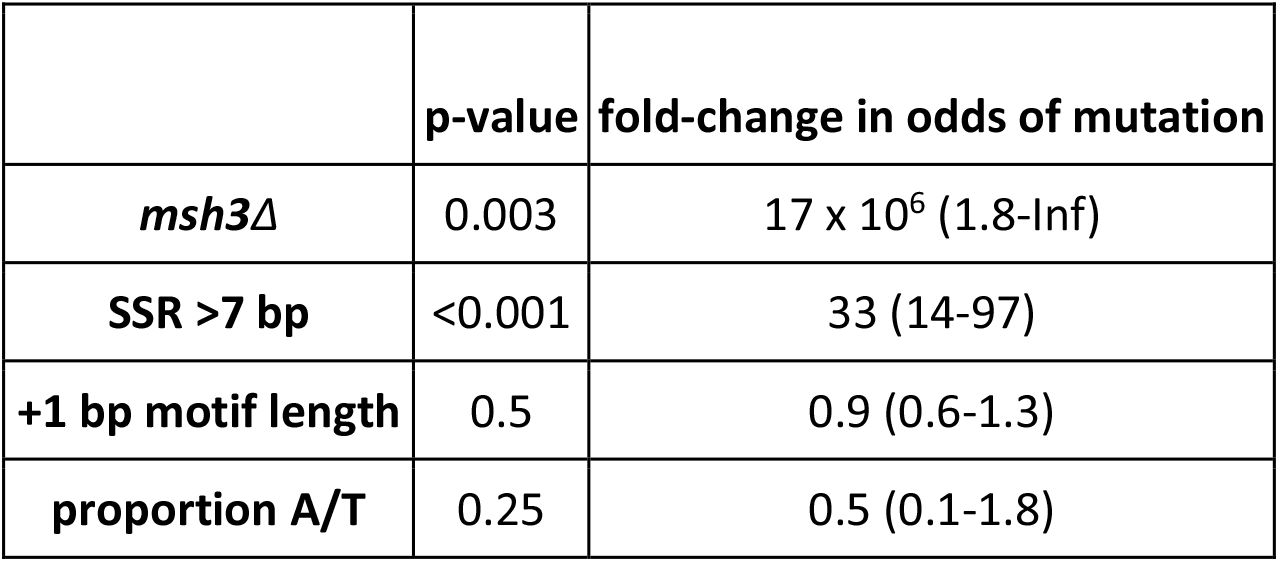
Effects of strain and SSR locus properties on the odds of SSR locus mutation. Fold changes in the odds of mutation, and p-values of fold change being significantly different than one, for *MSH3* genotype and various properties of SSR loci. The estimates of fold-change in odds of mutation are provided with a 95% confidence interval. Note that because no SSR mutations are found in *MSH3*^*+*^ strains, the coefficient on the effect of deleting *MSH3* (*msh3*Δ) is arbitrarily large, and has no upper bound.

Previous work showed that errors occurring in short SSRs are most often corrected through a polymerase-dependent mechanism, rather than by the MMR system (Lujan *et al*. 2015). To account for the possibility of additional error correction in short SSR loci, we included a parameter in our model for whether an SSR was shorter than 8 base pairs. We find that there is a 33-fold difference in per-base pair mutation odds between short and long SSRs (95% CI: 14-97-fold, p < 0.001) (**Table 1**).

The overall mutation rate in *msh3*Δ strains corresponded to ∼1.8 mutations/strain/200 generations. Although a total of more than 700,000 SSR loci passed filtration in *MSH3*^*+*^ strains, none of these threshold-passing SSRs were mutated in these strains. We therefore do not have enough data to calculate an estimate for the true SSR mutation rate in wild-type yeast, but we can calculate a lower bound on the fold-difference between *MSH3*^*+*^ and *msh3*Δ mutation rates: the 95% CI bound for the *msh3* mutation effect is a 1.8-fold increase in mutation rate in *msh3*Δ relative to *MSH3*^*+*^.

### SSR mutations are biased towards deletions

We next looked at the spectrum of mutations accumulated in SSR loci by the *msh3*Δ strains. Previous work has shown that SSR mutations display a bias towards deletions in homopolymers (Lang *et al*. 2013; Lujan *et al*. 2015). We observed a similar mutation pattern in the absence of *MSH3* (**Figure 3A**). In addition, we also observed a bias towards deletions in dinucleotide repeats and trinucleotide repeats, with most identified mutations being deletions of a single motif copy. We identified insertions among all three common categories of repeats. The increase in the rate of deletions relative to insertions was only significant among homopolymers (p = 0.001, 0.5, and 0.113 for homopolymer, dinucleotide, and trinucleotide repeats, respectively; no mutations passed thresholding in tetranucleotide repeats). Finally, Lang *et al*. identified an increase in the rate of substitutions in the proximity of SSRs in *msh2*Δ MA strains (Lang *et al*. 2013). We identified 4 substitutions within filtration-passing SSR regions in *msh3*Δ MA strains, at an overall rate of ∼1 substitution per 40,000 bp per generation. This is about 1.8x the non-SSR substitution rate; however, this difference is not statistically significant (p = 0.3).

**Figure 3:**
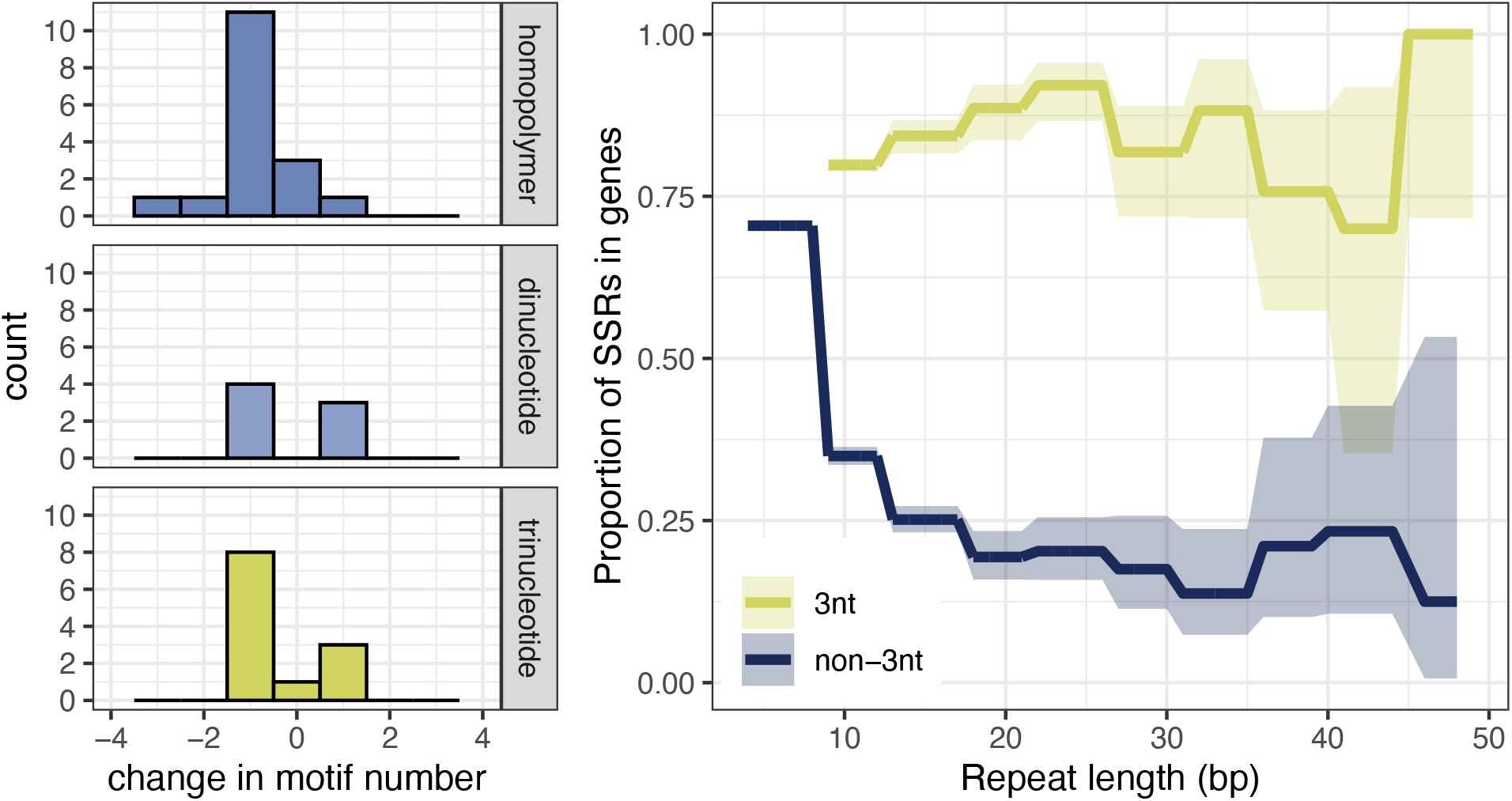
Spectrum of SSR mutations in *msh3*Δ strains. (A) Rates of insertions and deletions across motif lengths. In every SSR category (except tetranucleotide, where no mutations passed filtration), deletions were more frequent than insertions, with single-motif deletions being the most common category. Single-nucleotide substitutions inside SSRs are shown as having a change in motif number of 0. (B) Proportion of trinucleotide and non-trinucleotide SSRs in coding regions of the genome, as a function of total repeat length.

We also looked at the genomic locations of the accumulated SSR mutations. In particular, we looked at whether the mutations fell within or outside gene bodies. Seventy-three percent of the *S. cerevisiae* genome is genic and, in line with this, 69% of the SSR loci in our reference list fall within genes. This proportion, however, is dependent on motif length and locus length (**Figure 3B**). Trinucleotide repeats are found more frequently within genes (80%) than non-trinucleotide repeats (69%) (Fisher’s exact *P* < .001). Loci consisting of 10 or more repeats are less frequent than expected within genes (Fisher’s exact *P* < .001), although this dependency on length is only observable for non-trinucleotide repeats. The observed bias against non-trinucleotide repeats, especially long repeats, within gene bodies indicates the presence of selective pressure against potential frameshift mutations in coding regions, which would be highly deleterious (Metzgar *et al*. 2000).

Because selection should be minimal during an MA experiment, we expected the proportion of SSR mutations falling within genes to align with the previously described observations, with ∼70% of mutations falling within genes. Instead, we observed that SSR mutations were less likely than expected to occur within gene regions, with only 44% (16/36) falling within genes (Fisher’s exact *P* = .005). A similar bias was reported by Zhu et al. (2014): the authors observed a bias against indels, but not substitutions, within gene regions. We also tested whether this bias was influenced by motif length, and found that it was present for non-trinucleotide mutations (Fisher’s exact P < .001) but not for trinucleotide mutations (Fisher’s exact P = 0.9). This result would suggest that a fraction of spontaneously occurring indels are deleterious enough to not be observed in the final MA lines, even under reduced selection. However, as mentioned above, the proportion of non-trinucleotide SSRs in genes depends on the total length of the repeat, and on average only ∼25% of non-trinucleotide SSRs longer than 10 bp fall within genes (**Figure 3B**); this is close to the proportion of mutations in non-trinucleotide SSRs found in genic regions (6 out of 21). Because loci longer than 8 bp are the most likely to mutate (as shown in the previous section), this dependency of the SSR genic proportion on repeat length likely explains the observed bias against indels in genic regions.

## Discussion

Understanding the function of the Mismatch Repair Complex is critical for understanding the mutational processes that drive evolution and many aspects of human disease. We explored the role of the MutSβ MMR complex by accumulating mutations in a set of budding yeast lines mutant for the *MSH3* gene. We measured the genome-wide substitution rate in these strains, and found that this rate is not significantly affected by MutSβ function; our substitution rate estimate of 3.5 × 10^−10^ mutations/bp/generation is very close to a previous estimate of haploid mutation rate in yeast (4 × 10^−10^ mutations/bp/generation) (Sharp *et al*. 2018). We also devised a filtering strategy for SSR mutation calls that allows for accurate estimation of the SSR mutation rate despite inherent differences in sequencing accuracy among different classes of SSRs. Using this method, we found that deleting *MSH3* results in a >1.8-fold increase in the mutation rate of SSRs. We showed that deletion of *MSH3* results in a high rate of single-motif deletions, and (less frequently) insertions, in SSR loci. Finally, we showed that the mutation rate in SSRs at least 8 base pairs long is dramatically increased in *msh3*Δ mutants relative to shorter SSRs, lending further support to a model in which indels occurring within a few base pairs of the polymerase active site can be repaired by the polymerase, rather than by MMR machinery (Lujan *et al*. 2015).

A key difficulty in calling mutations in SSRs is the correlation between *in vivo* locus mutability and sequencing error, combined with the high overall error rate in sequencing SSR loci. This relationship is especially problematic when the total number of mutations in the sequenced strains is low, as the amount of SSR mutation signal is overwhelmed by the sequencing error noise. Previous studies have taken one of two general approaches to address this issue. Setting stringent requirements that all reads for a locus support a single genotype (e.g. (Lang *et al*. 2013; Haye and Gammie 2015), which is possible in haploid strains, can minimize the number of false positive calls when combined with a high threshold for the minimum number of reads per locus. However, as we have shown, when this approach is used to estimate the relative mutation rates of different SSR categories, it likely biases estimates of mutation rates against the most frequently mutated categories of loci, which are more likely to have reads supporting incorrect alleles. Unlike other types of sequence-inference error, increasing sequence coverage does not help with this bias. Indeed, the severity of the bias likely increases with increasing read depth, as observing more reads increases the chance that a locus is excluded from analysis due to a single mismatching read. Another approach has been to calibrate error rates specific to each SSR (Gymrek *et al*. 2012; Willems *et al*. 2017). Methods using this approach allow genotyping loci with divergent reads and work in diploids, but require a large amount of sequencing data, and may result in effectively overfitting locus-specific error rates. The latter issue is especially problematic when the total number of true mutations is low, as even a small amount of incorrect high-confidence calls can significantly skew mutation rates.

In this work, we use a different approach: rather than fitting a model to each individual locus, we use confidence calls from across tens or hundreds of similar loci to set different confidence thresholds for loci with similar properties. Pooling confidence scores across large groups of similar loci decreases the chance of overfitting and spurious false positives while also minimizing bias in mutation calls against more frequently mutated loci. When true mutation numbers are low, as in the present study, our method requires setting stringent thresholds that result in high false negative rates. We estimate a false negative rate of 40% among *msh3*Δ strains in the present study; it is difficult to estimate the false negative rates in other studies for comparison. However, we show that our method produces consistent mutation rate estimates across a wide range of filtration thresholds, effectively sacrificing the ability to call individual SSR mutations for an accurate estimate of the overall mutation rate.

We urge caution in estimating the rates of SSR variants, especially when these occur rarely (such as in mutation accumulation experiments). In addition to the issues described above, read length is likely a key determinant of bias in SSR mutation calls, with loci less likely to be called accurately as their length approaches the length of each read. Furthermore, accurate estimation of mutation rates in SSRs likely requires high read depth: in our study, threshold-passing SSR loci were sequenced with a median sequencing depth of 62x. Although the strategy we describe allows for mitigation of biases in SSR mutation rate estimates, it cannot compensate for insufficient read depth or read length. Importantly, the strategy described here also cannot be used directly for calling mutations in diploids, as heterozygous and homozygous calls inherently have different confidence (ΔGL) levels, and most mutations are heterozygous.

We estimate the SSR mutation rate in the absence of the mismatch repair gene *MSH3* to be ∼1 mutation/haploid genome/110 generations. Every indel observed in this study was in an SSR locus, and nearly every indel removed or inserted a single copy of the repeated SSR motif; the only exceptions to this were two homopolymers in which two and three of the SSR nucleotides were removed, respectively, and a single trinucleotide repeat in which a non-motif triplet was deleted from the middle of the SSR locus in a single strain. These observations are consistent with previous studies of SSR mutation rate (Kunkel and Erie 2015). We also observe a higher rate of deletions compared to insertions in SSR loci, which has been shown to be the case in homopolymers but not di- or tri-nucleotide repeats in *msh2*Δ mutants, e.g. (Lang *et al*. 2013; Lujan *et al*. 2015). We also observed indels in 6- and 10-bp repeats in *msh3*Δ lines, although the number of these mutations was too small to determine whether their occurrence in mismatch repair-deficient lines occurred by chance or due to an inherent difference in the rate of long SSR mutations between *MSH3*^*+*^ and *msh3*Δ strains; similarly, Sia et al. (1997) reported an increase in the mutation rate of microsatellites with 8-bp repeats in *msh3*Δ mutants.

Previous work has suggested complex roles for the MutSα and MutSβ complexes. Abrogation of MutSα function seems to affect both substitution and indel rate (Kunkel and Erie 2015). Although defects in MutSβ function result primarily in defects in repair of short indels (Kunkel and Erie 2015), mutations in *MSH3* have been shown using reporter constructs to have an effect on the spectrum of substitution mutations, but not their rate (Harrington and Kolodner 2007; Lamb *et al*. 2022). We do not detect a statistically significant difference in the number of substitutions between *MSH3*^*+*^ and *msh3*Δ strains, although the total number of mutations in our study is likely too small to test for an effect of the *msh3*Δ mutation on the spectrum of substitutions. The spectrum of mutations within SSR loci in *msh3*Δ strains from this study, and that previously reported for *msh2*Δ strains (Lang *et al*. 2013; Lujan *et al*. 2015), are similar: most mutations are deletions of a single motif copy, with a smaller number of insertions of a single motif copy or deletions of multiple motif copies (in the case of homopolymers). Reporter-based studies in *msh2*Δ mutants have identified differences in the SSR mutation rate of A/T- and G/C-rich loci in yeast (Gragg *et al*. 2002); however, studies in *msh2*Δ mutation accumulation lines have been equivocal about the effect of A/T proportion on SSR mutation rate (Lang *et al*. 2013; Lujan *et al*. 2015). Although we do not observe a significant dependence of SSR mutation rate on A/T proportion, this may be the result of a relatively small sample size of mutations.

Importantly, our work confirms results from mutation accumulation in an *msh2*Δ background demonstrating that the mutation rate of short SSR loci (up to 7-10 bp) is not as strongly affected by abrogation of mismatch repair as the mutation rate of longer SSR loci is (Lujan *et al*. 2015). This is not simply a result of fewer opportunities for mutation in shorter loci, as we find that the *per-base pair* odds of mutation of an SSR locus are ∼30-fold lower in loci shorter than eight base pairs long. Lujan *at al*. have previously shown that in the *msh2*Δ background, the rate of mutation for these short loci can be increased by abrogating the repair capabilities of DNA polymerase (Lujan *et al*. 2015). Our work therefore supports a model in which the probability of an indel-causing mismatch occuring is dependent on the length of an SSR, but such mismatches are efficiently repaired by polymerase in short SSRs; in longer SSRs, mismatch repair requires the MutSβ complex, and deleting members of this complex significantly increases the mutation rate for long SSR loci specifically.

## Materials and Methods

### Strains

The parent strain of the MA experiments is a haploid line derived from a single spore of the MA ancestor described in (Joseph and Hall 2004; Hall *et al*. 2008), with genotype *ade2, lys2-801, his3-ΔD200, leu2–3*.*112, ura3–52, ho*. A single colony derived from this ancestor founded strain *s*.*EP049*, which is the *MSH3*^*+*^ ancestral strain used in this study.

To construct the *msh3*Δ ancestor, *s*.*EP049* was transformed with a linear construct containing homology upstream and downstream of the *MSH3* gene flanking a Hygromycin/5-fluorodeoxyuridine positive/negative selection cassette (Alexander *et al*. 2014) that was flanked by two 50-bp internal homology sites; spontaneous recombination between these sites results in the excision of the selection cassette, leaving behind a single copy of the internal homology and a *Cyc1* terminator sequence. *msh3*Δ transformants were selected on hygromycin, genotyped, and re-selected on 50 μg/mL 5-fluorodeoxyuridine to select for strains with the selection cassette removed. The resulting *msh3Δ::Cyc1T* strain, founded by a single colony, was designated *s*.*EP060*.*3* (see **Supplementary Table 1** for a complete list of strains).

### Mutation accumulation

To perform mutation accumulation, single YPD-grown colonies of *s*.*EP049* and *s*.*EP060*.*3* were re-streaked on YPD; each resulting colony founded a single MA line, with five *MSH3*^*+*^ lines and 36 *msh3*Δ lines. Respiring *ade2* mutant colonies have a pink tint after two days of growth on YPD, developing a distinct red color after an additional two days. As in (Joseph and Hall 2004; Hall *et al*. 2008), we used this color difference to ensure selection of respiring non-petites during mutation accumulation; to facilitate this visual discrimination, a single *MSH3*^*+*^ petite line was passed through mutation accumulation alongside the others as a reference. Each transfer was performed in duplicate. To avoid unconscious bias in the subculture procedure, the two pink colonies that were closest to a pre-marked spot on the plate were chosen at every passage.

Colonies of each line were transferred every two days, and both colonies that were re-streaked were frozen in 50% YPD, 15% glycerol. The two colonies from each line used at each transfer were designated as ‘primary’ and ‘secondary’, and only the ‘primary’ re-streak of each line was used in the following transfer, except in cases when it turned out to be a petite, in which case a colony from the ‘secondary’ streak was used. A single line (*A17*) was petite in both the primary and secondary transfer near the last generation, and was removed from further analysis; one additional line (*A10*) was removed due to potential contamination during final MA generations. After each transfer, a mixture of many yeast colonies from the transfer plate was also frozen in 50% YPD, 15% glycerol. Transfers were halted for ∼3 months after the first 6 transfers (∼120 generations) due to an interruption in lab activities caused by the COVID-19 pandemic; mutation accumulation was resumed by streaking frozen whole colonies. A total of 10 transfers (∼200 generations of mutation accumulation) were performed.

### Sequencing

Cultures derived from single frozen colonies from the final MA transfer, as well as from ancestral MA strains, were grown in SC media and DNA extraction was performed as in (Schwartz and Sherlock 2016). Nextera library preparation was performed as in (Baym *et al*. 2015), but with 14 PCR cycles instead of 13. Bead cleanup was modified to optimize selection of 500-600 bp fragments: libraries were initially incubated with 0.53x volume AmpPure beads. Supernatant was saved, beads were washed with water, and then supernatant was incubated with original beads + 0.1x original volume AmpPure beads, followed by washing beads with 75% Ethanol and elution of DNA in 10 mM Tris pH 8, 1 mM EDTA, 0.05% Tween-20.

All sequences were deposited in SRA under project PRJNA871948, with run numbers listed in **Supplementary Table 1**. Confirmatory Sanger sequencing was performed using primers listed in **Supplementary Table 6**.

### Mutation calling with FreeBayes and filtration

Mutations were called using a custom pipeline written in nextflow (Di Tommaso *et al*. 2017), which is available at https://github.com/Siegallab/ssr_mutation_rate_pipeline. In short, reads were aligned using *bwa mem* (v0.7.17) (Li 2013) to a reference genome derived by updating the *S. cerevisiae S288C* reference genome with the ‘ancestral’ MA mutations identified in (Zhu *et al*. 2014). Duplicate reads were removed using GATK (v4.1.9) (Auwera and O’Connor 2020), and alignment files for libraries from the same strain were merged together using Picard (v2.17.11) (Broad Institute 2018).

To identify SSR loci, we first ran Tandem Repeat Finder (TRF) (Benson 1999) v4.09 with suggested parameters, except minimum alignment score, which was set to 3. TRF fails to identify a large number of short SSRs; therefore, we also performed a string search genome-wide for homopolymers with 4-10 repeats, and di- and tri-nucleotides with 3-10 repeats. The results of this search were joined with TRF results; repeats were filtered as suggested in (Willems 2017). Finally, any overlapping loci with non-identical motifs were split in such a way as to maximize the combined alignment score of the two motifs. The complete list of 274,411 SSR loci used in this study can be found in **Supplementary Table 7**.

We initially ran FreeBayes (v1.3.4) (Garrison and Marth 2012) with the default parameters, except *min_mapping_quality* was set to 1; any calls with a *QUAL* value greater than 1 were used in downstream analysis. This created a list of calls at sites, including SSRs, where at least one strain was mutated relative to the reference. In order to get call likelihood values for non-mutant SSR loci, any loci called as mutant were removed from the full list of SSRs. The SSR list was then converted to a vcf, with the alternate allele at each locus listed as a missing value.

FreeBayes was then re-run with the vcf of non-mutant SSR loci provided as *--variant-input*, and with the *--only-use-input-alleles, --min-alternate-count 0, --min-alternate-fraction 0*, and *--min-coverage 0* options. In this mode (and in the absence of a provided alternate allele), FreeBayes evaluated the likelihood of each unmutated SSR locus as compared to a version of the locus one motif repeat shorter than the original. The resulting calls and likelihood values were joined with the list of calls from the initial round of FreeBayes analysis.

In order to filter out spurious mutation calls, we removed any calls identified within 100 base pairs of a telomere, centromere, or LTR transposon (**Supplementary Table 8**); we also removed any calls inside the rDNA-containing regions of chromosome XII, and in the mitochondria. Only calls sequenced with a read depth of at least 10x were included. We also removed a small number of mutations falling in non-SSR repetitive regions and having low call confidence (differences <20 between log-likelihood of top calls). Finally, non-SSR mutations found in multiple strains were removed, as explained in *Results*.

### Mutation calling with muver

In addition to calling SNMs with FreeBayes, we performed SNM calling for all MA strains using muver, a tool specifically designed for the analysis of MA experiment data (Burkholder *et al*. 2018). Muver’s pipeline includes an alignment step, performed with Bowtie2 (version 2.3.5), and a variant calling step, during which SNMs and small indels are identified using GATK (version 3.8). Muver allows the user to indicate the ancestor strain of the experiment and, in a final step, calls substitutions and small indel mutations occurring in the MA strains compared to the ancestor. We ran muver on the *MSH3*^*+*^ and *msh3*Δ strains separately, specifying s.EP049 and s.EP060.3 as the ancestor strain, respectively. We filtered muver’s results to exclude mutations called in low mappability regions of the genome, including centromeric and telomeric regions and LTRs. Mutations occurring on mitochondrial DNA were also filtered out. Among the *MSH3*^*+*^ strains, only C3 had 3 mutations compared to the ancestor, whereas muver identified 117 mutations across the 34 *msh3*Δ strains. 91 of the called mutations fall into loci that we classified in our analysis as SSRs leaving 26 non-SSR loci mutated in the *msh3*Δ strains. Of these, 2 mutations were absent in FreeBayes calls; both of these loci were Sanger sequenced and found to be unmutated.

### Mutation calling with MSIsensor

In addition to FreeBayes, mutations in SSR regions were identified with MSIsensor, a tool designed to identify microsatellite instability in paired tumor-normal sequence data (Jia *et al*. 2020). First, we ran the *scan* command to identify all SSR regions in the reference genome, using the following parameters: *-l 4, -m 50, -r 3, -s 4*. The resulting file was filtered to remove loci within LTRs, telomeres, centromeres, and rDNA repeats on Chrom XII, as well as mutations occurring on mitochondrial DNA, using bedtools (v2.29.2) (Quinlan and Hall 2010). We then ran the *msi* command on all possible ancestor-MA strain pairs for both the *MSH3*^*+*^ and *msh3*Δ strains, specifying the following parameters: *-c 15, -l 4, -p 4, -m 120, -q 3, -s 3, -w 120, -f 0*.*1*.

### Modeling SSR locus mutation probabilities

To model the probability of SSR locus mutation, we modeled the odds of the observed mutation rate as a function of a baseline mutation rate, the locus length in base pairs, and any coefficients of interest:

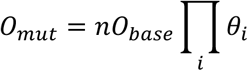

Where *n* is the length of the locus in base pairs, *O*_*base*_ is a ‘baseline’ per-basepair mutation rate, and each *θ*_*i*_ is a coefficient by which a given parameter *i* affects the odds of mutation.

Modeling was performed with a binomial generalized linear model in R (R Core Team 2020). Because no SSR mutations were found in *MSH3*^*+*^ strains, the effects of SSR parameters on mutation rate were modeled using data from *msh3*Δ strains only (this should have little or no effect on model parameters, but significantly improves model behavior). A model including only a boolean parameter representing long (>7 bp) vs short SSRs (in addition to locus length in base pairs and baseline mutation rate) was used as a null model for calculating the significance of A/T proportion and motif length on SSR mutation rate; to calculate the significance of long vs short SSRs, this model was compared to a null model in which the odds of mutation are determined only by a baseline mutation rate and the length of the locus in basepairs. Finally, to calculate the significance and confidence intervals for the effect of the *msh3*Δ mutation, we ran the model above on the full dataset, including coefficients for long vs short SSRs (which had the only significant effect on SSR mutation rate) and *MSH3* mutation status.

All confidence intervals were calculated using likelihood profiling.

## Acknowledgements

We thank the NYU Genomics Core for help with sequencing, and the Hittinger lab and Alison Gammie for sharing materials. This work was supported in part through the NYU IT High Performance Computing resources, services, and staff expertise. This work was supported by National Institutes of Health grant R35GM118170 (to MLS). RFS is a Professor at the Cancer Research Center Cologne Essen (CCCE) funded by the Ministry of Culture and Science of the State of North Rhine-Westphalia.

