## Supplemental Figures and Table Legends for "The rate of spontaneous mutations in yeast deficient for MutSβ function"

### Supplementary Tables

**Supplementary Table 1**
Mutation accumulation and corresponding SRA identifiers

**Supplementary Table 2**
SNMs identified in MA strains outside of SSR loci; note that the two mutations in strain *A4* on chromosome XI are within 19 bp of each other, and therefore counted as part of a single locus.

**Supplementary Table 3**
% genome that is above 10x coverage in *both* the MA strain listed and its ancestor, and which is not part of a repeat (telomeric, centromeric, or LTR regions)

**Supplementary Table 4**
Insertions/deletions identified in MA strains; ancestral sequence listed on top, MA line sequence on the bottom

**Supplementary Table 5**
Mutations in SSR loci; ancestral sequence listed on top, MA line sequence on the bottom. A single SSR locus is found in two strains. One mutation (Chrom VII:675527-675544) is likely to be a false positive: it is borderline significant, and the same mutation is called in many other strains at $\Delta$GL below the $\Delta$GL quantile threshold we implemented.

**Supplementary Table 6**
Primers used for amplifying the *MSH3* knockout cassette and genotyping. For genotyping primers, the F primer of the pair was typically used for Sanger sequencing the PCR product.

**Supplementary Table 7**
List of SSR loci used in this study

**Supplementary Table 8**
List of genomic regions excluded in this study due to being part of LTR, telomere, centromere, or ribosomal array (not including the 100-bp buffer used in calling mutations)

#### Supplementary Figures

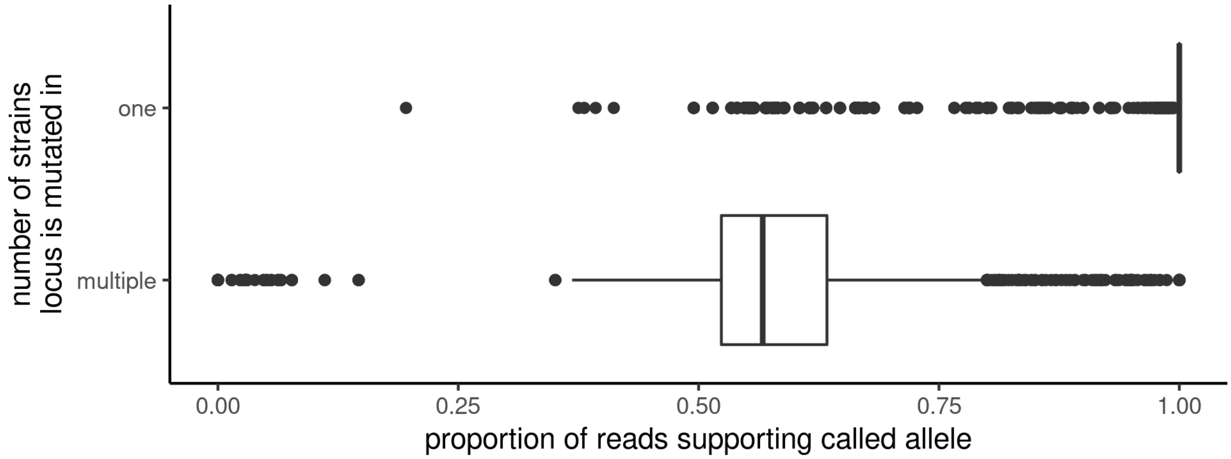


**Supplementary Figure 1**
Allele calls for loci mutated in multiple strains have a low proportion of reads supporting them

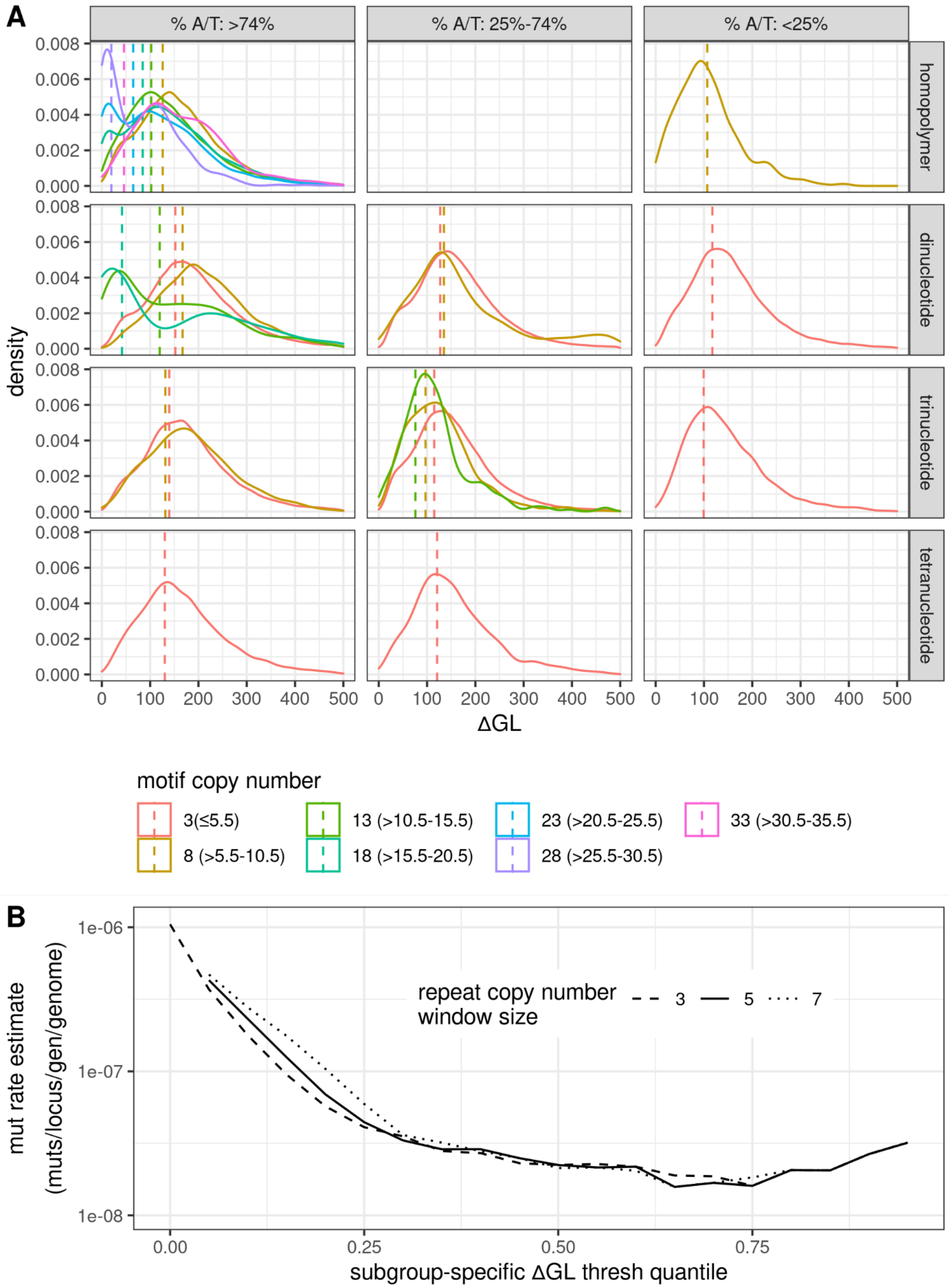
A
% A/T: >74%
% A/T: 25%-74%
% A/T: <25%
homopolymer
dinucleotide
trinucleotide
tetranucleotide
density
ΔGL
motif copy number
3(≤5.5)
8 (>5.5-10.5)
13 (>10.5-15.5)
18 (>15.5-20.5)
23 (>20.5-25.5)
28 (>25.5-30.5)
33 (>30.5-35.5)
B
mut rate estimate
(muts/locus/gen/genome)
repeat copy number
window size
3
5
7
subgroup-specific ΔGL thresh quantile

**Supplementary Figure 2: Filtering of SSR locus calls**

(A) Empirical distributions of ∆G values across SSR locus properties. For representation clarity, thresholds for only a subset of motif copy numbers are shown. Dotted line in each plot represents the 35th-percentile cutoff ∆G value for each category; distributions include ∆G values for all SSR calls within the repeat copy number window specified in parentheses. Locus call ∆G values vary across locus properties, but setting a within-group ∆G percentile-based threshold ensures that mutations in loci that have generally lower call confidence can still be identified.

(B) Choice of grouping window size for repeat copy numbers during quantile-based ∆GL filtration does not substantially affect the estimated mutation rate. Solid line corresponds to **Figure 2C.**

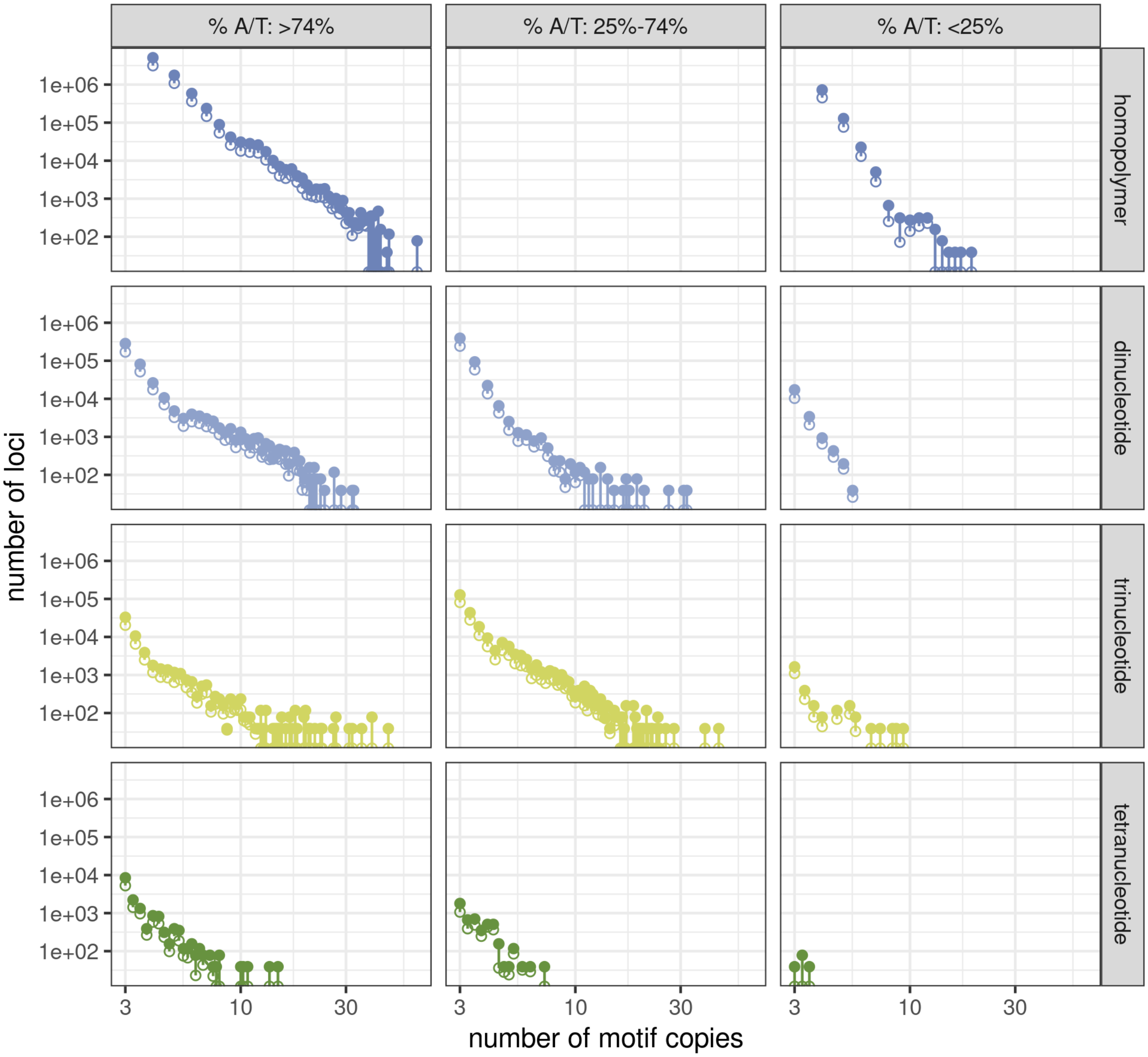


**Supplementary Figure 3: Filtering of SSR locus calls**

Removing rare groups of loci in filtration does not significantly bias the types of loci genotyped by motif complexity or A/T proportion. Solid circles represent the total number of potential calls in the dataset (locus number x number of MA strains) corresponding to a motif complexity, A/T proportion, and copy number; empty circles represent the total number of calls made after filtration. With the exception of long SSR loci (>35 base pairs), which are rare in the yeast genome, a consistent proportion of potential calls in each category passed filtration.
